# Synthetic type III-E CRISPR-Cas effectors for programmable RNA-targeting

**DOI:** 10.1101/2024.02.23.581838

**Authors:** Daniel J. Brogan, Elena Dalla Benetta, Tianqi Wang, Calvin P. Lin, Fangying Chen, Harry Li, Claire Lin, Elizabeth A. Komives, Omar S. Akbari

## Abstract

The recent discovery of the type III-E CRISPR-Cas effector class has reshaped our fundamental understanding of CRISPR-Cas evolution and classification. Type III-E effectors are composed of several Cas7-like domains and a single Cas11-like domain naturally fused together to create a single polypeptide capable of programmably targeting and degrading RNA. Here we identify a novel composition of a type III-E-like effector composed of Cas7-like and a Cas1-like domain, that can be engineered into an active chimeric RNA-targeting Cas effector and presents a new function of Cas1 in RNA-targeting. Furthermore, we demonstrate a unique modularity of type III-E effectors by methodically substituting domains between orthogonal type III-E proteins to engineer compact synthetic Cas effectors. We refine our methods to generate several compact effectors for programmable RNA-targeting in mammalian cells. Cas7-S represents a new understanding of type III-E architecture and modularity, and provides a platform for engineering genome engineering technologies from the blueprint of nature.

## Introduction

The continuous evolution of CRISPR-Cas systems has resulted in the discovery and engineering of revolutionary genome engineering technologies. Recently, the type III-E CRISPR-Cas system was discovered (*1*) and characterized (*2*, *3*) and has changed how we understand Cas protein evolution. Type III-E effectors are categorized as class 1 CRISPR-Cas systems, which typically utilize a multiprotein complex to accomplish DNA- and/or RNA-targeting (*1*). However, type III-E Cas effectors accomplish programmable RNA-targeting by employing a single, large protein composed of four Cas7-like domains, one Cas11 domain, and a large insertion (INS) domain, hence being named Cas7-11(*4*, *5*) (or gRAMP for giant Repeat Associated Mysterious Proteid (*6*)). Recent studies have demonstrated that type III-E effectors are capable of programmable RNA knockdown in prokaryotic and eukaryotic cells, and do not exhibit RNA collateral cleavage activity that consistently limits applications of CRISPR-Cas13 systems (*7–10*). Using prokaryotic expression, it was demonstrated that Cas7-11 target binding activates a downstream caspase activity that results in cell dormancy, or death, and appears to serve as the primary immune response for type III-E CRISPR-Cas systems (*6*, *11*, *12*). Therefore when type III-E effectors are expressed independently of their protease signaling pathway, they operate as highly specific ribonucleases and their unique architecture provides a pathway for further engineering.

It is important to consider effector size when engineering CRISPR-Cas systems, and efforts have been made to generate compact effectors for almost all therapeutically relevant CRISPR-Cas systems. Usually, Class I CRISPR-Cas systems are extremely challenging to encode in therapeutic vectors since they require co-expression of multiple independent proteins (*13*), which highlights the advantage of using Cas7-11 instead. However, type III-E effectors are the largest single-effector Cas protein subtype discovered to date (*2*, *3*), which is a limitation for therapeutic applications, such as AAV delivery. For RNA-targeting CRISPR-Cas systems, Cas13 effectors, such as the compact Cas13d subtype (*14*, *15*), are more feasible for packaging, however severely limited by collateral cleavage of ssRNA (*7*). Recently, a small Cas7-11 (Cas7-11S) protein was engineered through deletion of the INS domain and replacement with a GS-linker (*4*), however the unique architecture of Cas7-11 characterized in this study suggests a unique modularity of type III-E effectors that can be exploited for engineering.

We envisioned type III-E effectors as a natural fusion of modular proteins with interchangeable orthologous domains. We reasoned that the unique assembly of, essentially, individual proteins into a single polypeptide, provided evidence of a naturally occuring fusion system. We hypothesized that interchanging domains between orthologs may enhance certain functions, such as improving catalytic activity and experimentally test this in mammalian cells. Moreso, we identify an interesting architecture of a type III-E-like protein composed of Cas7-like domains and a Cas1-like domain. We use this novel composition as inspiration to engineer recently characterized type III-E effectors into synthetic Cas effectors that we term Cas7-S. We demonstrate a likely novel function of Cas1 in RNA-targeting, and provide a method for designing synthetic RNA-targeting Cas effectors. Together this study provides a unique perspective on CRISPR-Cas biology, as well as an engineering approach to produce compact synthetic RNA-targeting Cas effectors.

## Results

### Identification of a type III-E-like architecture containing a Cas1 domain

To begin engineering type III-E effectors, we initially conducted a bioinformatic search and alignment to create a domain library (**Table S1**) and identify any variants with possible novel architectures. In our initial search we obtained an unusual blast hit for a protein denoted as a Cas1 nuclease (MBU1487208.1). We analyzed the domain architecture of this protein using HHPred and found the protein to be composed of three Cas7-like domains and one Cas1 nuclease domain (**Figure S1A-E**). For simplicity, we term this protein Cas7-1. We then searched for possible crRNAs associated with Cas7-1 and obtained a 38nt consensus direct repeat (DR). We obtained secondary structure predicted folds of the forward and reverse directions of the consensus DR **(Figure S2A-B)**, where we found the reverse direction to contain a similar stem-loop structure to the DR of DiCas7-11 **(Figure S2B-C)**. We biochemically tested Cas7-1 targeted cleavage activity with 5’ and 3’ orientations of the 2 DR sequences **(Table S3)** and observed no cleavage activity **(Figure S2D)**.

Although Cas7-1 demonstrated no enzymatic activity, the protein aligns well to all known type III-E effectors and through further analysis we found interesting features of the protein. Cas7-1 appears to lack an INS domain – a domain found in all known type III-E effectors (*3*) (**Figure 1A**). We also found Cas7-1 contains a conserved catalytic residue that aligns to all known Cas7.3 domains, as well as zinc-finger motifs unique to type III-E effectors (*4*) (**Figure 1B-E**). Lastly, the Cas1 domain aligns to a Reverse Transcriptase-Cas1 (RT-Cas1) fusion protein commonly associated with type III systems of all characterized subtypes (III-A, III-B, III-C, and III-D) (**Figure 1F, File S1**). In these systems, the active Cas1 domain is required for adaptation of new spacers from interfering RNA (*16*). We therefore hypothesized that Cas1 may function by orienting the target substrate RNA for cleavage. However, from the alignments and structural prediction (**Figure 1G**), it appears Cas7-1 is missing a portion of the N-terminus, including a CRISPR RNA (crRNA) processing domain and a secondary catalytic residue, which likely explains the lack of activity (**File S2**).

**Figure 1.**
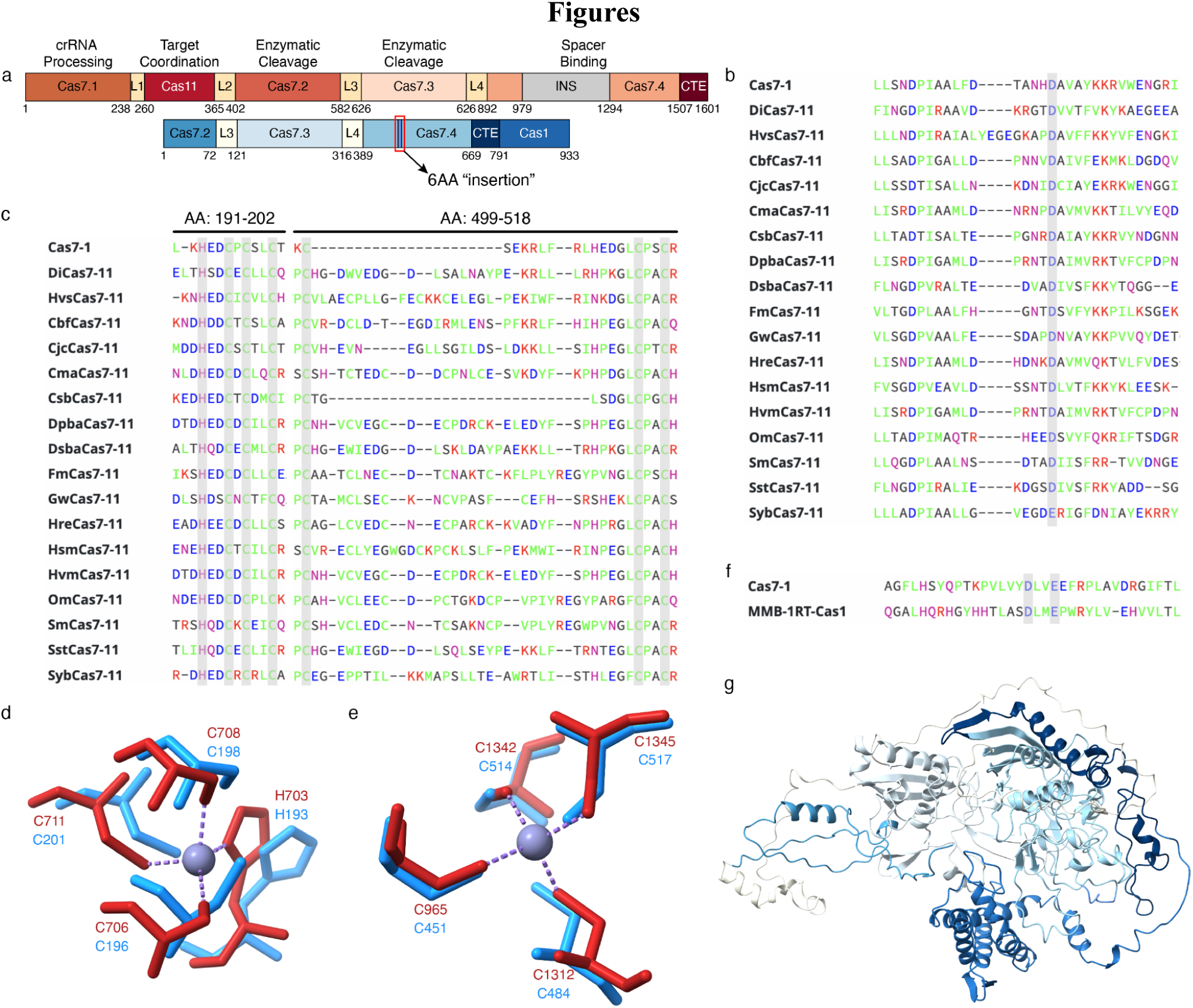
Cas7-1 sequence and predictive structure in relation to type III-E effectors. (**A**) Schematic representation of a typical Cas7-11 architecture (red) and the architecture of Cas7-1 (blue). (**B**) Alignment of catalytic residue in the Cas7.3 domain of Cas7-1 against published Cas7-11 proteins (*3*). (**C**) Alignment of zinc finger residues in the Cas7.3 and Cas7.4 domains of Cas7-1 and Cas7-11 proteins. Amino Acid (AA) position represents position in Cas7-1 protein sequence. (**D**) Structural alignment of Cas7.3 zinc finger residues of Cas7-1 (blue; H193/C196/C198/C201) with aligned residues of DiCas7-11 (red; H703/C706/C708/C711). (**E**) Structural alignment of Cas7.4 zinc finger residues of Cas7-1 (blue; C451/C484/C514/C517) with aligned residues of DiCas7-11 (red; C965/C1312/C1342/C1345). (**F**) Alignment of Cas1 domain of Cas7-1 to MMB1 RT-Cas1 fusion protein (NCBI Ascension here). Highlighted sections represent catalytic domains of Cas1, where E847 of Cas7-1 aligns to E870 of MMB1 RT-Cas1. (**G**) Predicted structure of Cas7-1 obtained from AlphaFold 2. Coloring of domains is as depicted in (**A**).

### Engineering synthetic Cas effectors with varying architectures

Based on prior characterization of type III-E effectors, we suspected the missing domains of Cas7-1 would abolish crRNA recognition and processing and, partially, cleavage activity. Therefore, we reasoned we could rescue activity of the Cas7-1 protein through the incorporation of Cas7.1 and Cas7.2 domains from *D. ishimotonii* Cas7-11 (DiCas7-11). We also hypothesized that these Cas7-S effectors could be generated using a method we term Orthologous Domain Substitution (ODS), where aligned domains from type III-E effectors are substituted between orthologs. To rescue Cas7-1 activity and test the ODS design method, we designed 16 initial Cas7-S effectors with different domain and linker substitutions, while simultaneously varying the overall architecture based on Cas7-1 or typical Cas7-11 proteins (**Figure 2A, Figure S3, Figure S4A, Table S2**). We also obtained predicted structures from AlphaFold (*17–19*) to align the models against the solved structure of DiCas7-11 and use them for downstream engineering (**File S3-25**).

**Figure 2.**
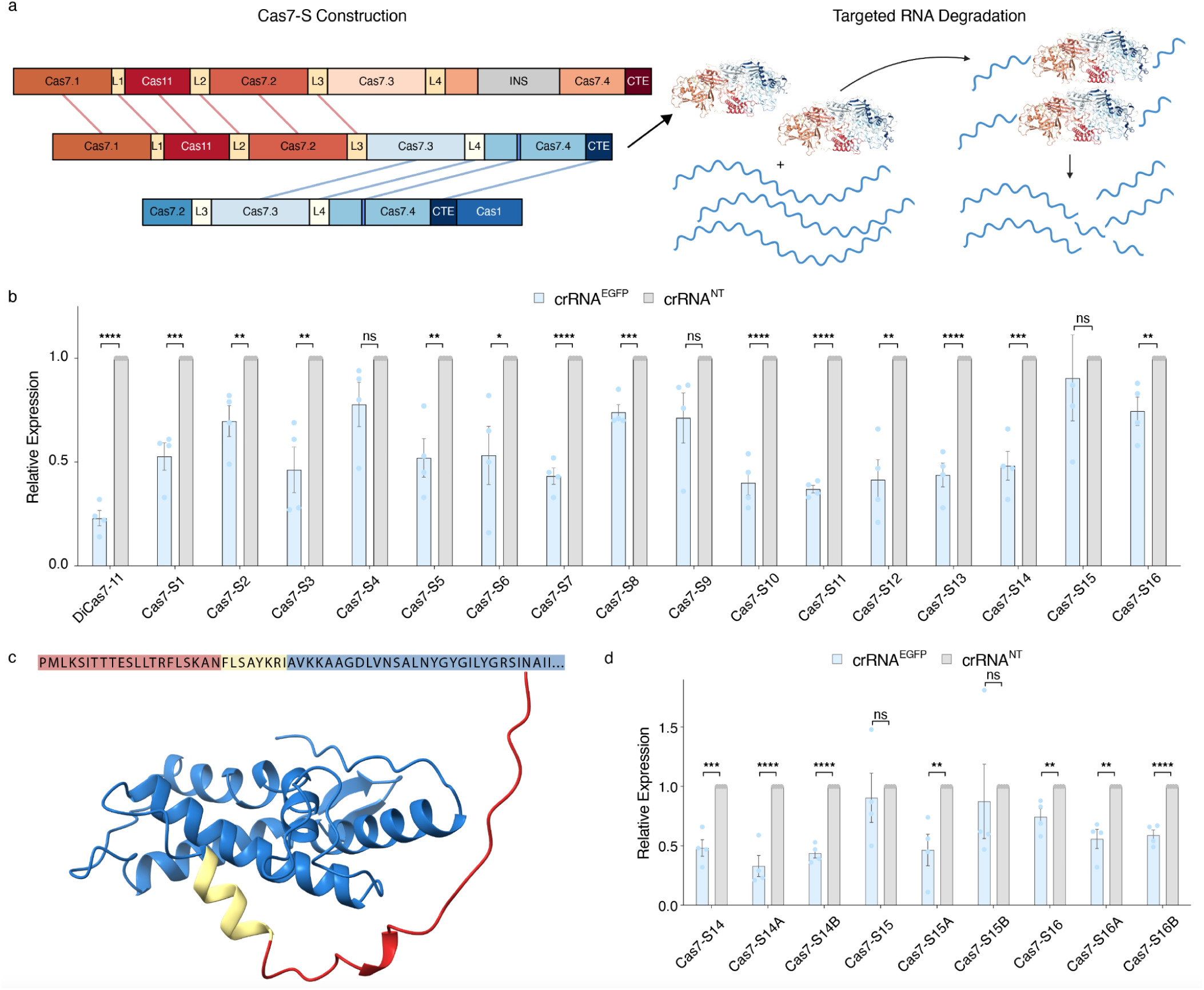
Cas7-S construction and analysis of initial variants. (**A**) Schematic representing the construction of Cas7-S10 and a depiction of RNA degradation by Cas7-S10. Red lines represent domains from DiCas7-11 and blue lines represent domains from Cas7-1. (**B**) qPCR analysis of EGFP targeted knockdown with Cas7-S variants that have a C-terminal orientation of the Cas1 domain and Cas7-S variants that orient the Cas11 or Cas1 domain internally in the typical Cas7-11 architecture. (**C**) Predicted structure and partial amino acid sequence of Cas1 domain with highlighted areas representing truncation of the domain. Red and yellow combined represent truncation for “A” variants, while red alone represents truncation for “B” variants. (**D**) qPCR analysis quantifying EGFP knockdown by Cas7-S variants with different truncations of Cas1. In all qPCR plots, significance is calculated and determined using unpaired t-test between crRNA^EGFP^ and crRNA^NT^. Error bars represent sem (n = 4).

Since no previous study has utilized a Cas1 domain in programmable RNA-targeting and we suspect the Cas1 domain would operate to orient the target substrate, we focused on Cas7-S variants that were either Cas1-based or Cas11-based (**Figure S3**). We divided the groups by architecture, aiming to determine if the Cas7-1 orientation is functional or the typical Cas7-11 architecture reigned supreme. We tested our Cas7-S variants in HEK293T cells using an EGFP reporter assay and analyzed samples via qPCR. crRNAs used in assessment contained DRs respective to DiCas7-11 and spacer sequences complementary to EGFP (**Table S3**). Although we observed significant RNA knockdown with the Cas7-1 architecture (Cas7-S1-9), most variants exhibited high variability and the best variant (Cas7-S7) includes the large INS domain (**Figure 2B**). In the Cas7-11 architecture group (Cas7-S10-16), we observed consistent RNA knockdown comparable to DiCas7-11 from multiple variants including one Cas1-based variant (Cas7-S14) (**Figure 2B**). We identified Cas7-S10-12 (where Cas7-S12 is nearly identical to Cas7-11s (*4*)) as the best Cas11-based effectors and Cas7-S14 as the best Cas1-based effector.

Due to the natural orientation of the Cas1 domain on the C-terminus of Cas7-1, we suspected the N-terminal region of the domain would be relatively unstructured and highly flexible in order to help orient the Cas1 domain underneath target substrates. We identified a 29AA stretch that aligned specifically to the N-terminus of the RT portion of RT-Cas1 fusion proteins (**Figure S5, File S26**). Using the predicted structure of Cas7-S14, we generated two truncated versions of the Cas1 domain (**Figure 2C**) and tested these truncations with Cas7-S14, Cas7-S15, and Cas7-S16. Aside from Cas7-S15B, we found both truncations generally improved knockdown activity for the Cas7-S variants and observed that complete deletion of the linker region consistently improved activity likely due to increased stabilization of the Cas1 domain (**Figure 2D**). Together these findings demonstrate RNA-targeting type III-E effectors are highly modular and Cas1 can be used in RNA-targeting applications.

### Programmable cleavage of ssRNA with Cas7-S effectors *in vitro*

With evidence of RNA knockdown in cell culture, we next wanted to characterize the mechanism of cleavage of the Cas7-S effectors. We focused on Cas7-S10 to demonstrate that substitution of enzymatic domains permits retention of enzymatic activity and that RNA-targeting is unperturbed. Since Cas7-S10 contains the Cas7.1 domain deriving from DiCas7-11, all crRNAs were designed with DRs respective to DiCas7-11 (**Table S3**). We ran an initial assay truncating the spacer of the crRNA and observed cleavage for spacers between 20-30nt in length. Interestingly, we noticed a substantial increase in both crRNA processing and target cleavage as spacers were shortened, with the best cleavage activity observed for 22-24nt spacers (**Figure S6A**). This increase in cleavage activity due to spacer truncation was further replicated with Cas7-S10 in cell culture via qPCR analysis (**Figure 3A**). We also extended incubation time of the cleavage reaction from 1.5 hours up to 6 hours. We found 1.5 hours of incubation is sufficient to obtain noticeable cleavage of the target substrate, however complete cleavage of the target substrate was observed after 6 hours of incubation (**Figure S6B**). This suggests Cas7-S10, like type III-E effectors, accomplish target cleavage slowly.

**Figure 3.**
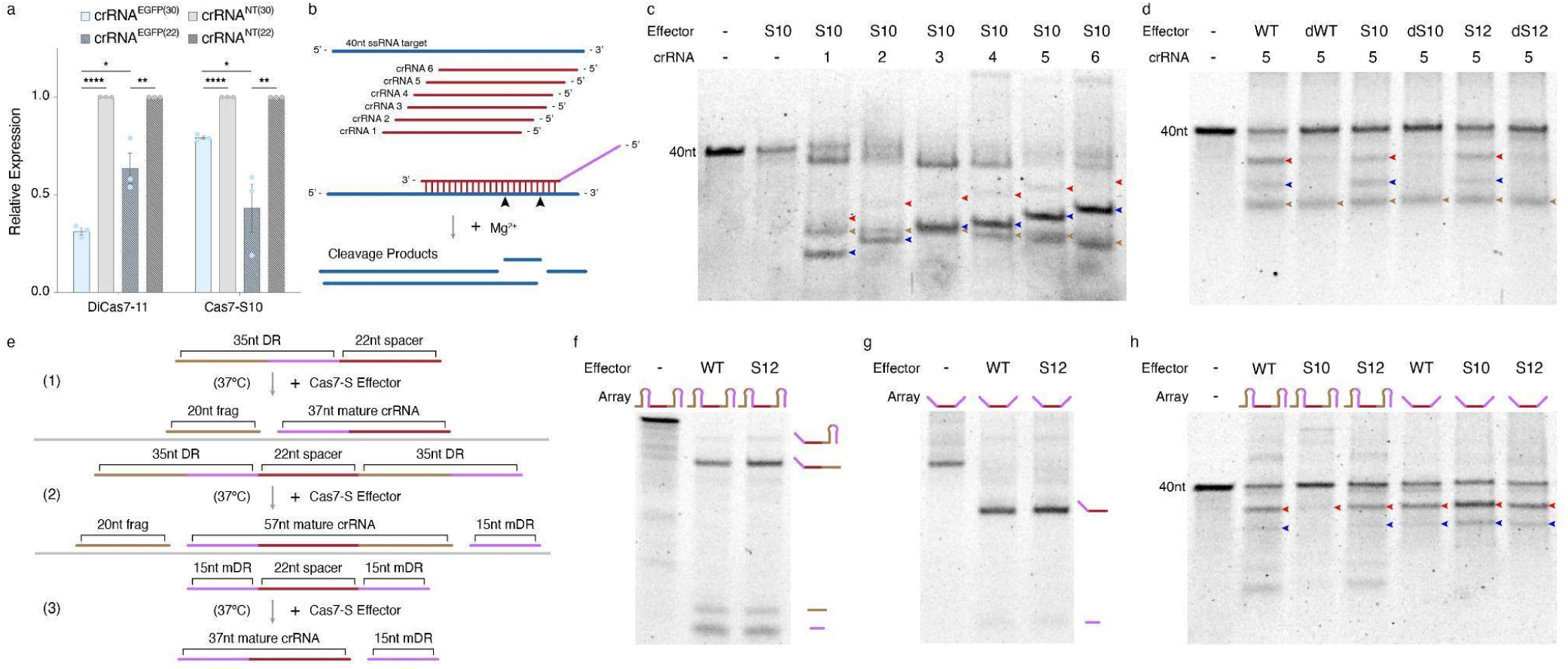
Biochemical assessment of Cas7-S cleavage activity. (**A**) qPCR analysis of EGFP knockdown comparing 30nt vs 22nt spacers for DiCas7-11 and Cas7-S10. Significance is calculated and determined using unpaired t-test between crRNA^EGFP^ and crRNA^NT^. Error bars represent sem (n = 3). (**B**) Schematic depicting the crRNAs used for assessment of *in vitro* cleavage activity and depiction of cleavage and expected cleavage products. Black arrowheads represent cut sites of crRNA-5. (**C**) Experiment tiling six 22nt crRNAs across the 40nt ssRNA target to obtain stepwise cleavage products. (**D**) Assessment of cleavage pattern and catalytic inactivation of DiCas7-11, Cas7-S10, and Cas7-S12 through site specific mutagenesis (D429A/D654A). (**E**) Schematic representation of different crRNA processing outcomes. (1) “single-guide” crRNA processing. (2) native array crRNA processing. (3) modified array crRNA processing. (**F-G**) crRNA array processing activity by DiCas7-11 (WT) and ΔINS mutant, Cas7-S12. Array structures represented by schematics above each lane. crRNA processing products and approximate location on gel depicted by schematics on the side of gel. (H) Comparison cleavage assay between WT DiCas7-11, Cas7-S10, and Cas7-S12 (ΔINS) with varying array structures. Red arrows represent cleavage by Cas7.2 only. Blue arrows represent cleavage products by Cas7.3 or from both Cas7.2/Cas7.3 active sites cleaving the target. Brown arrows represent 20nt processed DR fragments. Purple represents 15nt mature DR (3’ only).

To visualize both crRNA processing and target cleavage, we designed several crRNAs across a 40nt ssRNA target and incubated the Cas7-S effectors with unprocessed crRNAs (**Figure 6B**). Tiling the crRNAs across the ssRNA target and incubating for 6 hours to clearly visualize the complete cleavage product, we observed stepwise cleavage products demonstrating activity of both enzymatic domains (**Figure 3C**). Cleavage activity was completely abolished through mutagenesis of both active sites for DiCas7-11, Cas7-S10, and Cas7-S12 (D429A/D654A), while crRNA processing was unaffected (**Figure 3D**).

We investigated the crRNA processing activity further by introducing a full array structure containing the same spacer as crRNA-5 (**Figure 3E, Table S3**). Differing from other reports (*2*, *3*), we found that neither Cas7-S12, nor wild-type (WT) DiCas7-11, could completely process an array structure into a mature 37nt crRNA, but instead yielded an incompletely processed 57nt product (**Figure 3F**). We found only the Cas7.1 domain processed the DR producing a product where a 15nt mature DR (mDR) is upstream of the spacer and a 20nt DR fragment (DRf) remains at the 3’ end (**Figure 3E**). This suggests that type III-E effectors do not cleave off the DRf downstream of the spacer when processing arrays. However, by modifying the array structure to incorporate mDRs surrounding the spacer sequence, we were able to obtain a crRNA without excess nucleotides on the 3’ end (**Figure 3G**). This array design is extremely important for targeting and multiplexing applications with Cas7-S effectors, as we found Cas7-S10, which contains Cas7.3/Cas7.4/CTE domains from Cas7-1, RNA cleavage activity is partially inhibited when the DRf is present (**Figure 3H**). Together, these results suggest that Cas7-S effectors accomplish RNA cleavage like other Cas7-11 proteins and type III-E CRISPR-Cas systems use a particular crRNA structure and processing mechanism similar to previous reports of type I and type IV CRISPR-Cas systems (*20*).

### Generation of compact Cas7-S effectors for RNA-targeting

Cas7-11 proteins are the largest single-effector Cas protein family discovered (*2*, *3*). In therapeutic applications, this large size is a major obstacle for delivery modalities and efforts have quickly been made to reduce the size of these enzymes (*4*). We hypothesized the ODS design method could be applied to engineer compact effectors. To test this, we selected three effectors (DiCas7-11, HvsCas7-11, and Cas7-1) to generate a set of 36 compact Cas7-S effectors and further refine our ODS design method for systematic interrogation of domain arrangement (**Figure 4A, Figure S7, Figure S4B, File S27-S62**). We included Cas7.4 from Cas7.1 in all compact Cas7-S effectors and also maintained all linkers from DiCas7-11, except when HvsCas7.1 and HvsCas11 were encoded together. The compact effectors varied in size from 1233-1292AA in length and are far smaller than the average size of Cas7-11 proteins (>1600AA) (**Table S2**).

**Figure 4.**
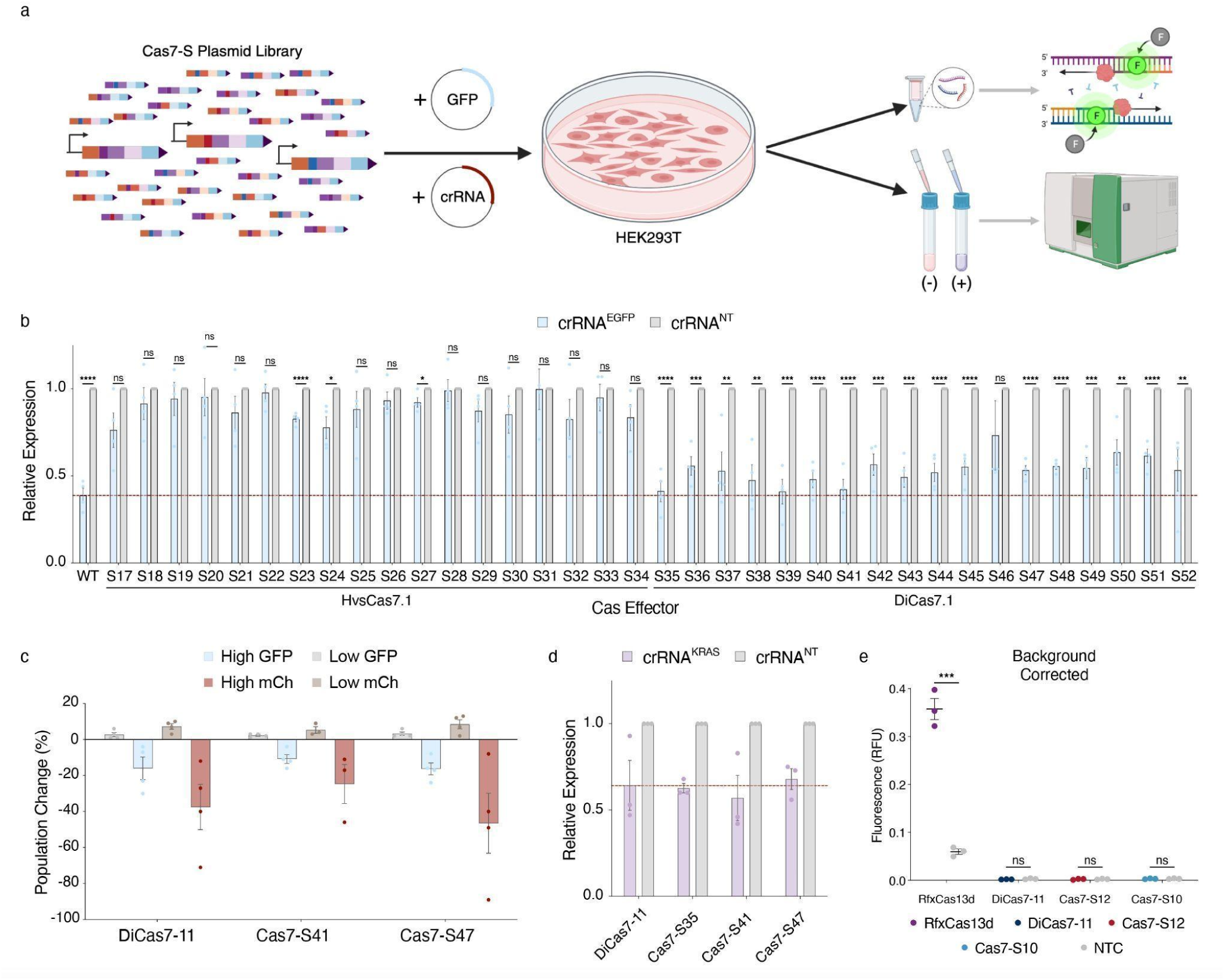
Generation and validation of compact Cas7-S effectors. (**A**) Schematic depicting workflow for analysis of Cas7-S library with refined ODS design method. (**B**) qPCR analysis of EGFP knockdown with all 36 compact Cas7-S effectors. Data is broken into two groups HvsCas7.1-based (S17-S34) and DiCas7.1-based (S35-S52) Cas7-S effectors. Dashed red line represents average knockdown by DiCas7-11 (WT). Significance is calculated and determined using unpaired t-test between crRNA^EGFP^ and crRNA^NT^. WT = wild type DiCas7-11. Error bars represent sem (n = 4). (**C**) Flow cytometry analysis of select Cas7-S effectors demonstrating effect of RNA knockdown on translation of EGFP and mCherry. Data depicts population change in High or Low GFP, or mCherry, expression groups (see methods for gating). Error bars represent sem (n = 4). (**D**) RNA knockdown of KRAS gene with DiCas7-11 and select Cas7-S effectors analyzed using qPCR. Error bars represent sem (n = 3). (**E**) SENSR assay determining ssRNA collateral activity of Cas7-S effectors depicted by background corrected fluorescence levels. RfxCas13d was used as a positive control. Error bars represent sem (n = 3). RFU = relative fluorescence unit.

To assess the RNA knockdown capabilities of the compact Cas7-S effectors, we analyzed these proteins using the previously described reporter assay in HEK293T cells, and quantified knockdown using qPCR and flow cytometry (**Figure 4A**). We found that Cas7-S effectors containing the DiCas7.1 domain resulted in consistent EGFP knockdown comparable to DiCas7-11, while effectors containing the HvsCas7.1 domain demonstrated little-to-no knockdown activity (**Figure 4B**). Within the DiCas7.1 groups, we also found that either DiCas11- or HvsCas11-based variants greatly outperformed Cas7-S effectors with a Cas1 domain, contrary to prior analysis of the first 16 variants, where Cas7-S14A demonstrated comparable knockdown capabilities to DiCas7-11. Of the Cas11-based effectors, Cas7-S35, Cas7-S39, and Cas7-S41 demonstrated knockdown comparable to DiCas7-11, where Cas7-S41 is the most compact (1241AA) and represents the smallest type III-E effector developed (**Figure S4C**). Interestingly, composing the Cas7-S effector of HvsCas7.2 and HvsCas7.3 resulted in high RNA knockdown regardless of the Cas11/Cas1 domain used, which was not observed for any other Cas7.2-Cas7.3 combination (**Figure 4B, Figure S4D**).

We further analyzed the Cas7-S effectors to gauge their ability to reduce the expression of target proteins. Using flow cytometry, we observed knockdown of EGFP and mCherry with Cas7-S41 and Cas7-S47 at comparable levels to DiCas7-11 for both targets tested (**Figure 4C**). This demonstrates that Cas7-S effectors can elicit phenotypic effects on a target of interest regardless of domain composition and at levels similar to wild-type type III-E effectors. Furthermore, we demonstrate Cas7-S effectors are capable of knocking down endogenous genes as effectively as DiCas7-11 (**Figure 4D**). We also show Cas7-S effectors do not possess the collateral cleavage activity found in other CRISPR-based RNA-targeting systems, such as Cas13. We accomplished this by running SENSR reactions (*21*) with Cas7-S effectors and comparing them to RfxCas13d. From the fluorescence assays, we found no evidence of collateral cleavage activity with any effectors composed of type III-E domains (**Figure 4E**), consistent with previous reports (*3*). Here we demonstrate that the Cas7-11 proteins are modular and can be engineered into compact, efficient, and highly specific Cas7-S effectors for RNA-targeting applications.

## Discussion

As new discoveries of CRISPR-Cas systems emerge, the continuous evolution of these enzymes reveals new possibilities for generating new genome engineering technologies. Here we identify a novel polypeptide composition similar to that of Cas7-11 proteins, termed Cas7-1, where the Cas1 domain appears to operate similarly to the Cas11 domain. Cas7-1 is also the first example of a type III-like effector naturally lacking an INS domain and demonstrating the possibility of a naturally compact type III-E effector. Using Cas7-1 and the Cas7-11 architecture, we developed an engineering method (ODS) to generate synthetic type III-E effectors capable of RNA-targeting. The ODS method reveals a unique modularity of the type III-E effector class, where orthologous domains are often interchangeable and minimally impact RNA knockdown activity. Furthermore, we demonstrate the applicability of this design method for generating the most compact type III-E effector to date with further improvements and applications possible.

Along with presenting an engineering method that manipulates the modularity of Cas7-11 proteins, we also uncover a unique crRNA processing activity where the mature crRNA contains a spacer surrounded by fragments of the DR on both the 5’ and 3’ end. This mode of crRNA processing has not been reported for type III-E or other single-effector CRISPR-Cas systems capable of array processing (*22*, *23*), and is reminiscent of crRNA processing by multiprotein type I CRISPR-Cas systems (*20*). For Cas7-S effectors, we found this crRNA processing activity to directly impact both array processing and targeting activity and we provide a solution for crRNA array design when applying Cas7-S effectors for RNA-targeting applications using a mDR array structure. These findings demonstrate the need for further exploration and characterization of Cas7-11 proteins, especially in the context of downstream engineering.

The ODS method and Cas7-S effectors are useful for protein and transcriptome engineering, respectively. Although we observed clear evidence of RNA-targeting *in vitro* and in mammalian cells, the level of activity exhibited by Cas7-S effectors in mammalian cells did not generally outperform WT DiCas7-11. This activity gap suggests further refinement is required to improve engineering of Cas7-S effectors. This refinement can be accomplished with larger libraries of Cas7-S effectors, incorporating machine learning to guide library design, or employing other downstream protein engineering methods, such as direct evolution (*24*, *25*). RNA cleavage by Cas7-11 enzymes is inherently slow due to the evolution as a caspase system for antiviral defense, which operates more effectively with longer target binding (*6*, *11*, *12*). This slow turnover is exemplified when comparing RNA knockdown by DiCas7-11 against RfxCas13d (**Figure S8**). Therefore, future studies should focus on increasing the rate of catalysis to enhance activity, while also maintaining high target specificity.

## Methods

### Bioinformatics

Cas7-1 was identified through BLAST queries of type III-E CRISPR-Cas effectors (MBU1487208.1). The amino acid sequence of Cas7-1 was submitted to HHPred for individual domain assignment (https://toolkit.tuebingen.mpg.de/tools/hhpred). Assumed DRs of Cas7-1 were identified by submitting metagenomic data related to Cas7-1 to CRISPR-Cas++ (https://crisprcas.i2bc.paris-saclay.fr/). DR secondary structure prediction was obtained using UNAFold (http://www.unafold.org/). Amino acid sequence alignments were performed using the Clustal Omega.

### Design and cloning of constructs

Plasmids for expression of effectors in HEK293T cells were designed with a CMV promoter and bovine growth hormone (bGH) terminator. Effector plasmids were generated using standard gibson assembly methods. crRNA plasmids were designed for expression driven by a U6 promoter and terminator. crRNA expression plasmids were generated using standard golden gate assembly methods. Select plasmids are available at https://www.addgene.org/ (**Table S4**).

### Mammalian cell culture

HEK293T cell lines were obtained from the American Type Culture Collection (ATCC CRL-3216) and used in all mammalian cell experiments. HEK293T cells were cultured in Dulbecco’s Modified Eagle Medium (DMEM) (Thermo Fisher Scientific 11995073) supplemented with 10% Fetal Bovine Serum (FBS) (Corning 35-011-CV) and 1% penicillin-streptomycin (Thermo Fisher Scientific 15070063). Transient transfections were carried out with Lipofectamine 3000 (Thermo Fisher Scientific L3000001) according to the manufacturer’s protocol.

### RNA-knockdown Assay

To assess RNA knockdown, HEK293T cells were co-transfected with 20ng of either the EGFP or mCherry reporter plasmid, 600ng of a respective Cas effector plasmid, and 300ng of crRNA plasmid. Cells were seeded in 48-wells 18-20 hours prior to transfection at seeding densities of 32,000 cells per well. Following transfection, cells were incubated for 48 hours at 37°C, 5% CO_2_. After 48 hours, cells were removed from the plate with 250µL of RNAprotect Cell Reagent (Qiagen 76104) and stored at -20°C for at least 24 hours prior to extraction. For each effector tested, four biological replicates were performed and a non-target crRNA was used as a control.

### Total RNA collection and qPCR

To measure the reduction of endogenous or reporter genes, the transfected cells, preserved in RNAprotect, were lysed using QIAshredder (Qiagen 79656). Total RNA was then extracted using the RNeasy Mini Kit (Qiagen 74106) according to the manufacturer’s protocol. Following extraction, the total RNA was treated with DNase (Thermo Fisher Scientific AM1907) to remove any remaining DNA. Sample RNA concentration was analyzed using a Nanodrop OneC UV-vis spectrophotometer (Thermo Fisher Scientific NDONEC-W). Approximately 0.5μg of total RNA was used to synthesize cDNA with a RevertAid First Strand cDNA Synthesis Kit (Thermo Fisher Scientific K1622). cDNA was diluted (1:1000) for qPCR analysis. qPCR was performed with SYBR green (qPCRBIO SyGreen Blue Mix Separate-ROX 17-507B, Genesee Scientific), using 4μl of dilute cDNA in each 20μl reaction containing a final primer concentration of 200nM and 10μl of SYBR green buffer solution. Three technical replicates were performed for each reaction. Pipetting was performed using EpMotion® 5075 (Eppendorf). The following qPCR protocol was used on the LightCycler® 96 (Roche): 3 min of activation phase at 95°C, 40 cycles of 5 sec at 95°C, and 30 sec at 60°C. The relative expression levels of EGFP were calculated using the manufacturer’s software and the delta-delta Ct method (2–ΔΔCt), with the GAPDH gene serving as a reference (*26*). To assess differences in EGFP expression between the non-target and targeting crRNA conditions for each effector, statistical analysis was performed using GraphPad Prism 10. Specifically, data were analyzed using an unpaired t-test.

### Flow Cytometry

Fluorescence knockdown was assessed using flow cytometry on the BioRad S3e cell sorter system. To prepare cells for analysis, transfected HEK293T cells were washed with PBS pH 7.4 (Gibco 10010031), all media was removed, then 40µL of Accumax (VWR AM105) was added to the well. 230µL of cold PBS pH 7.4 was added to the wells and cells were resuspended and transferred to a cell strainer (Corning 352235). Cells were then assessed on the S3e cell sorter with a count of 80,000 cells per sample. Flow cytometry data were analyzed using Floreada.io (https://floreada.io/) (**Figure S9**).

### Recombinant Protein Expression and Purification

Plasmids used for protein expression in *E. coli* were generated by subcloning ORFs from plasmids used for mammalian cell culture expression, along with an N-terminal 6xHis-SUMO into a pET28a vector backbone. Plasmids were then transformed into Rosetta™ 2(DE3)pLysS Competent Cells (Novagen). Single colonies were used to inoculate a 40mL LB overnight starter culture. The starter cultures were then transferred to a 1L LB culture in 4L baffled flasks and grown at 37°C with shaking (200 rpm) until OD600 reached ∼1.0. Cultures were then brought to 4°C before protein expression was induced with 0.2 mM isopropyl β-D-thiogalactopyranoside (IPTG) and grown for 18-20 hours at 18°C. Cells were then pelleted and resuspended in a lysis buffer containing 50mM Tris-HCl pH 8.0, 300mM NaCl, 10mM imidazole, 5% glycerol (v/v), and 3 mM β-mercaptoethanol supplemented with Protease Inhibitor Cocktail (Sigma P2714), 5mM PMSF, and 2.5 U/mL salt active nuclease (Sigma SRE0015). Cells were lysed via sonication and clarified by centrifugation. His-tagged protein was bound to 2mL HisPur™ Ni-NTA Resin (Thermo 88221) equilibrated with lysis buffer in a glass Econo-Column® Chromatography Columns (Bio-Rad) by flowing through the clarified lysate. A wash step was then performed by adding 12 column volume (CV) of a wash buffer containing 50mM Tris-HCl pH 8.0, 500mM NaCl, 30mM imidazole, and 5% glycerol followed by elution with 7 CV elution buffer containing 50mM Tris-HCl pH 8.0, 300 mM NaCl, 300mM imidazole, and 5% glycerol. The 6xHis-SUMO tag was cleaved off by dialyzing the eluate overnight at 4°C with ∼0.6 mg of in-house produced 6xHis-tagged Ulp1 (Yeast SUMO Protease) in a dialysis buffer containing 20mM HEPES-NaOH pH 7.5, 250mM NaCl, 5% glycerol, and 1mM DTT. Dialyzed sample was then flowed over the same Ni-NTA column equilibrated in a buffer containing 20mM HEPES-NaOH pH 7.5, 250mM NaCl, 25mM imidazole, 5% glycerol, and 1mM DTT to remove additional *E. coli* impurities, Ulp1, and the 6xHis-SUMO tag. Cation exchange chromatography was performed with 2x1mL HiTrap Heparin HP Column (Cytiva) with a NaCl gradient from 200-1000mM NaCl followed by gel filtration chromatography with a HiLoad 16/600 Superdex 200 column equilibrated in 20mM HEPES-NaOH pH 7.5, 600mM NaCl, 5% glycerol (v/v), 2mM DTT on an ÄKTA Pure (Cytiva). Proteins were then concentrated to ∼10µM and stored in small aliquots at -80°C for future use.

### Nucleic acid target and crRNA preparation

Synthetic ssRNA templates and the full DiCas7-11 crRNA array were ordered custom from IDT. All other crRNAs were produced in house. To generate crRNAs, a dsDNA template with a T7 promoter incorporated was produced through templateless PCR and purified with MinElute PCR Purification Kit (Qiagen 28004). dsDNA templates were converted to ssRNA through *in vitro* transcription using MEGAScript^TM^ T7 Transcription Kit (Invitrogen AM1334) and purified with MEGAClear^TM^ Transcription Clean-Up Kit (Invitrogen AM1908).

### Nuclease assays

Nuclease assays were run in 10µL reactions with the following final concentrations: 500nM protein, 200nM crRNA, 240nM probe, 40U RNase Inhibitor, and 10mM MgCl_2_. Reactions were supplemented with a custom 10X reaction buffer (200mM HEPES (pH 7.5), 600mM NaCl) and incubated at 37°C. Times were varied based on experiment. Upon completion, reactions were denatured with 2X RNA dye (NEB B0363S) at 95°C for 10 min. 10µl of dyed samples were loaded onto a prerun (200V for 60 min) 15% TBE-Urea PAGE gel (BioRad 4566055) and resolved at 200V for 35 min. Gels were then stained with SYBR Gold (Invitrogen S11494) and incubated for 10 min at room temperature on a shaker and washed in 1X TBE for 10 min on a shaker. Stained gels were imaged using Enduro^TM^ GDS (Labnet). SENSR assays were conducted following a previously described protocod (*21*) with slight alterations: 100nM of RfxCas13d incubated in reaction and 200nM of DiCas7-11 or Cas7-S effector incubated in reaction.

## Data Availability

Select plasmids have been sent to Addgene and will be available for ordering. For other plasmids not found on Addgene, please submit a request to OSA.

## Acknowledgments

This work was supported by NIH/NIAID (R01GM132825-04, R21RAI149161A, and DP2AI152071) awarded to OSA. Thank you to the members of the Komor lab for advice in establishing systems for assay development and providing template plasmids essential to experimentation. We would also like to thank Kevin Corbett for his critical advice provided.

## Author Contributions

Conceptualization: DJB. Methodology: DJB, EDB, CPL. Investigation and analysis: DJB, EDB, TW, CPL, FC, HL, CL. Supervision: OSA, EAK. Writing (original draft): OSA, DJB, EDB, TW, CPL, EAK.

## Competing interests

DJB and OSA have a patent pending relating to this work. OSA is a founder of Agragene, Inc. and Synvect, Inc. with equity interest. The terms of this arrangement have been reviewed and approved by the University of California, San Diego in accordance with its conflict of interest policies.

---

**Figure S1.**
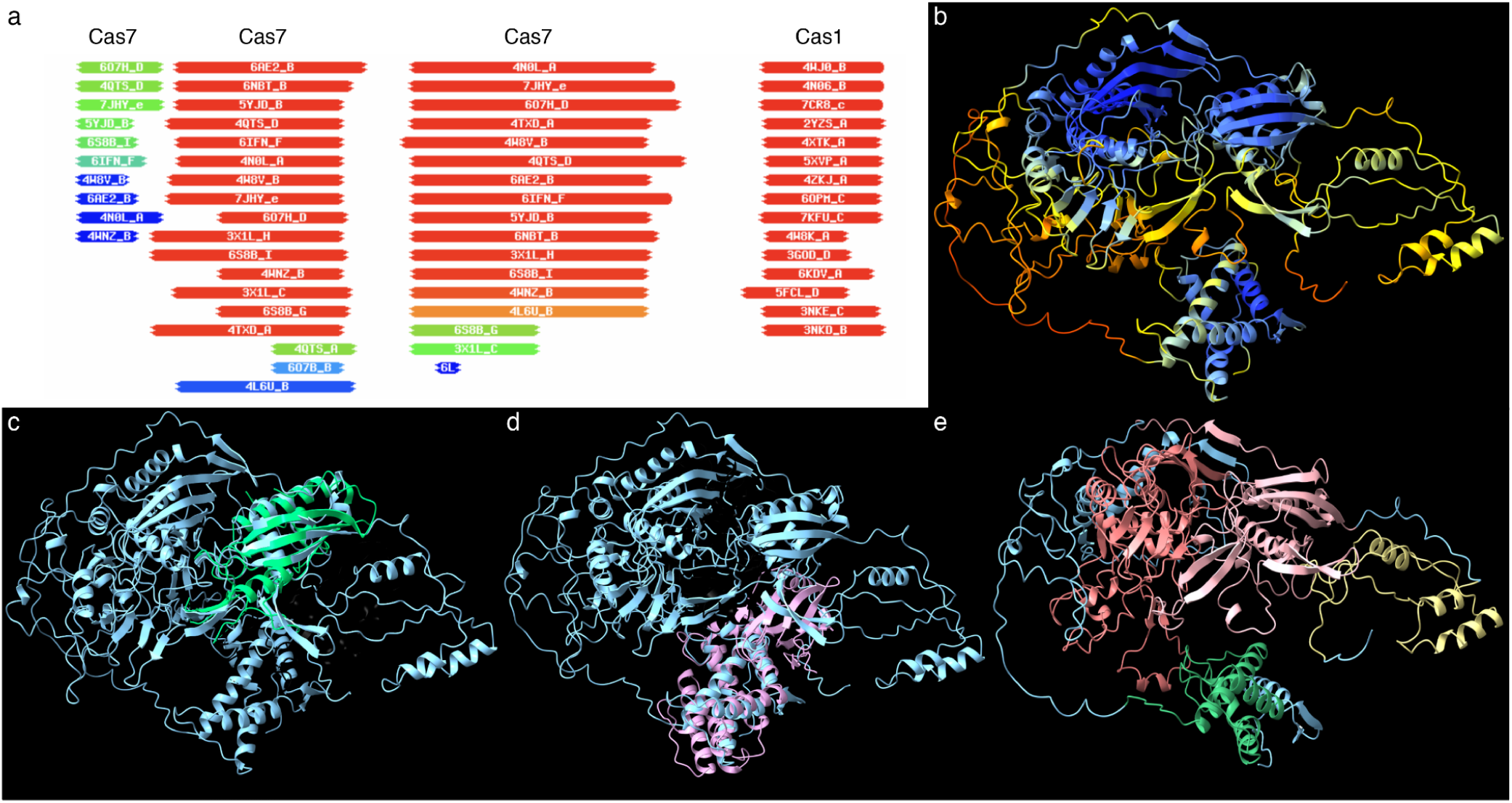
Sequence and structure prediction of Cas7-1. (**A**) HHpred domain assignment results. Domains are segmented based on alignments to most likely results. Results are depicted from red to blue (most likely to least likely). (**B**) AlphaFold predicted structure and heat map representation of Cas7-1. Coloring represents high confidence (blue) down to low confidence (red) of prediction. (**C**) Spatial alignment of Csm3 (green; PDB: 6nbt) over Cas7-1. (**D**) Spatial alignment of Cas1 (purple; PDB: 4w8k) over Cas7-1. (**E**) Recoloring of predicted domains from HHPred analysis. Khaki represents Csm3 domain 1 (mid confidence), light pink represents Csm3 domain 2 (high confidence), light coral represents Csm3 domain 3 (high confidence), and sea green represents Cas1 (high confidence).

**Figure S2.**
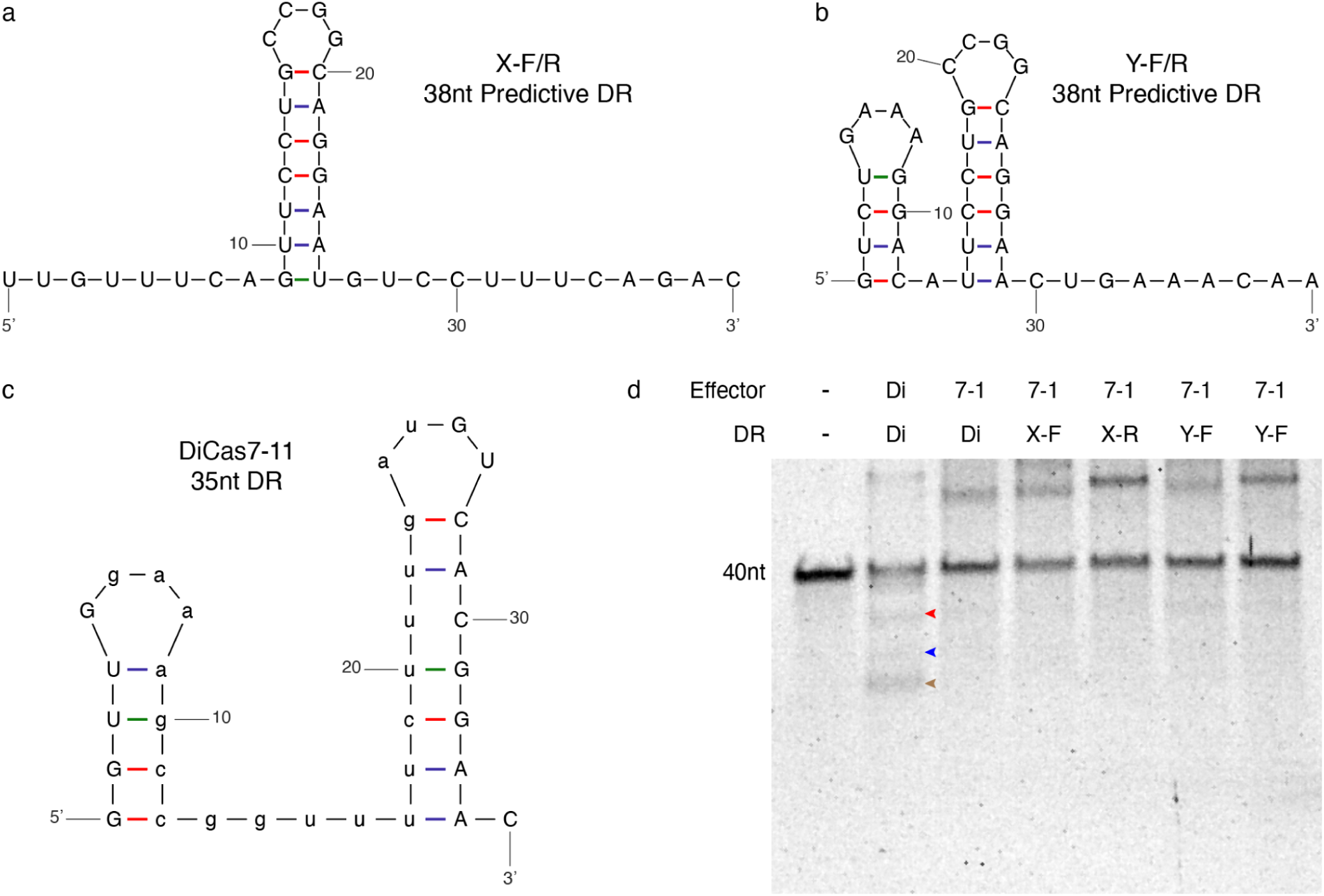
Cas7-1 predicted DR processing and cleavage assessment. (**A**) Predicted fold of predicted 38nt Cas7-1 direct repeat sequence DRX. (**B**) Predicted fold of predicted 38nt Cas7-1 direct repeat sequence DRY. F represents 5’ orientation and R represents 3’ orientation of DR related to the spacer. (**B**) Predicted fold of known 35nt DR for DiCas7-11. (**D**) Gel electrophoresis assessment of crRNA processing and targeted cleavage with predicted DRs of Cas7-1 compared with DiCas7-11. Red arrow indicates cleavage products from Cas7.2 domain, blue arrow indicates cleavage products from Cas7.3 and/or Cas7.2/Cas7.3 domain(s), and brown arrow indicates 20nt product from DR processing.

**Figure S3.**
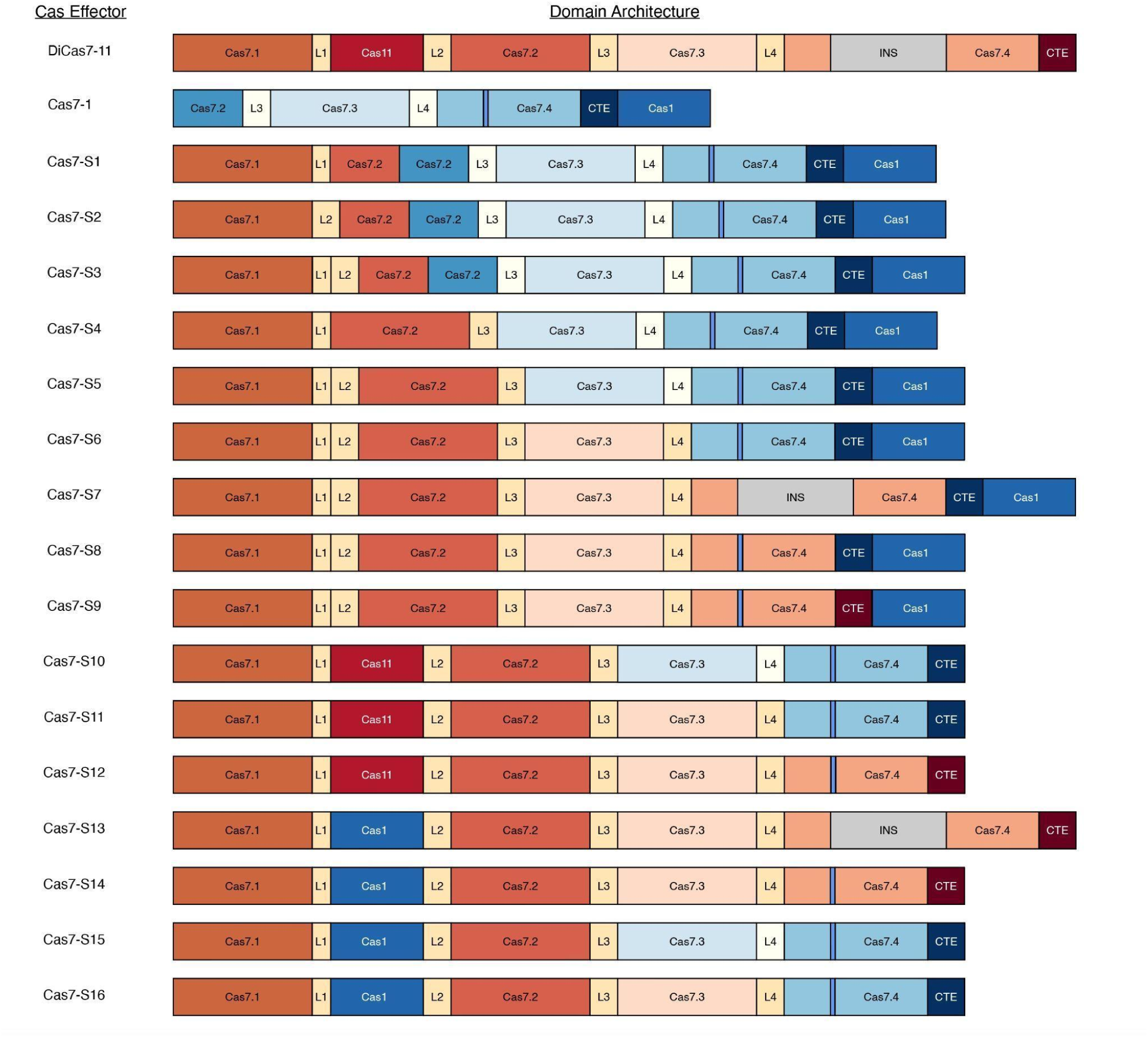
Domain breakdown of initial Cas7-S variants. Original proteins to build the initial Cas7-S variants are DiCas7-11 (red) and Cas7-1 (blue). Color scheme represents which native protein individual domains originate. Domains are not scaled.

**Figure S4.**
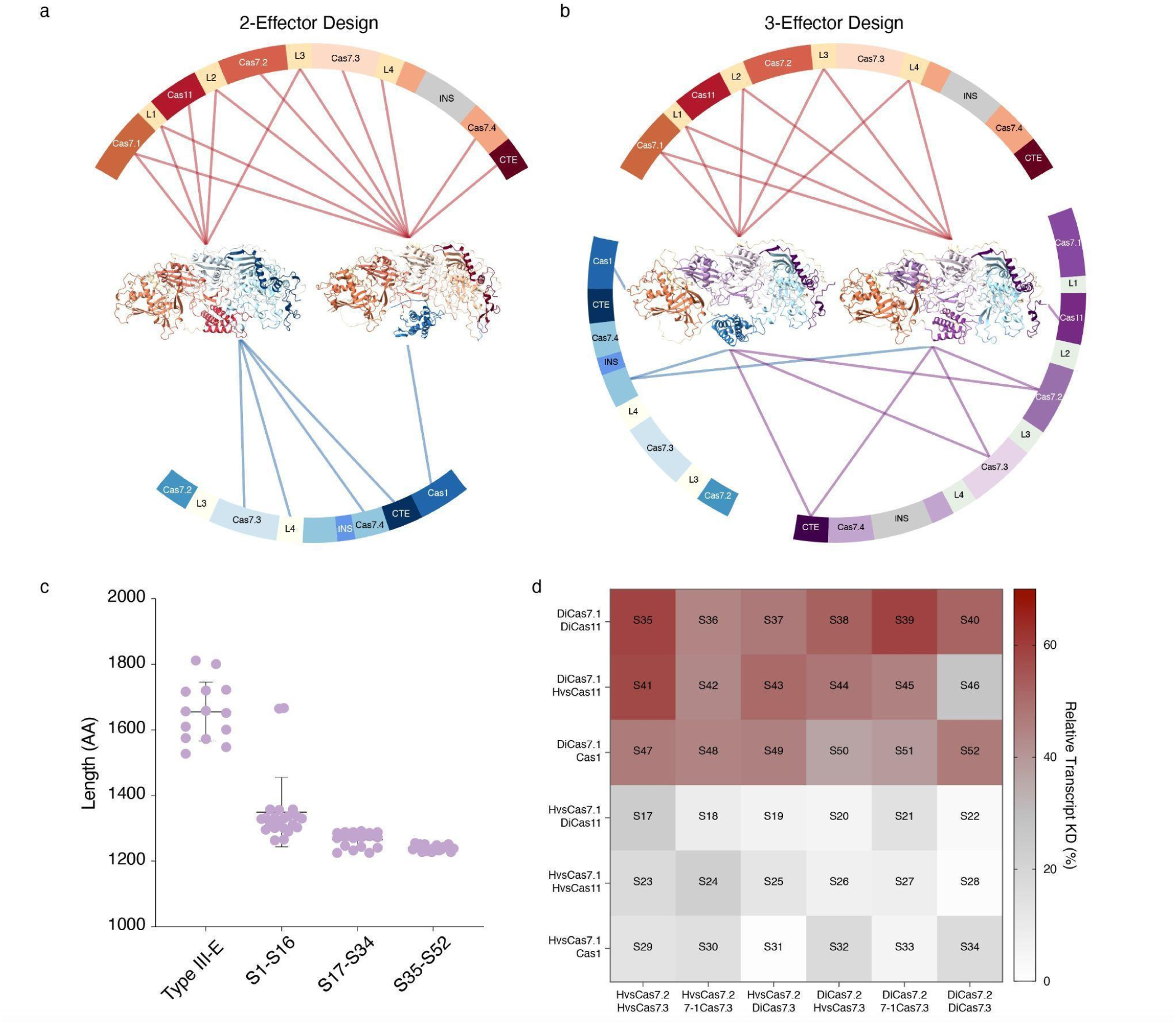
Design and assessment of Cas7-S effectors. (**A**) Schematic representation of 2-effector design strategy with Cas7-S10 (left) and Cas7-S14 (right) depicted. (**B**) Schematic representation of 3-effector design strategy with Cas7-S41 (right) and Cas7-S47 (left) depicted. (**C**) Size comparison between type III-E effectors and the various Cas7-S effectors generated in this study. (**D**) Heat map representation of qPCR data from Cas7-S17-52 analysis. The more intense the red saturation the stronger the knockdown level. 4 replicates in each cell.

**Figure S5.**
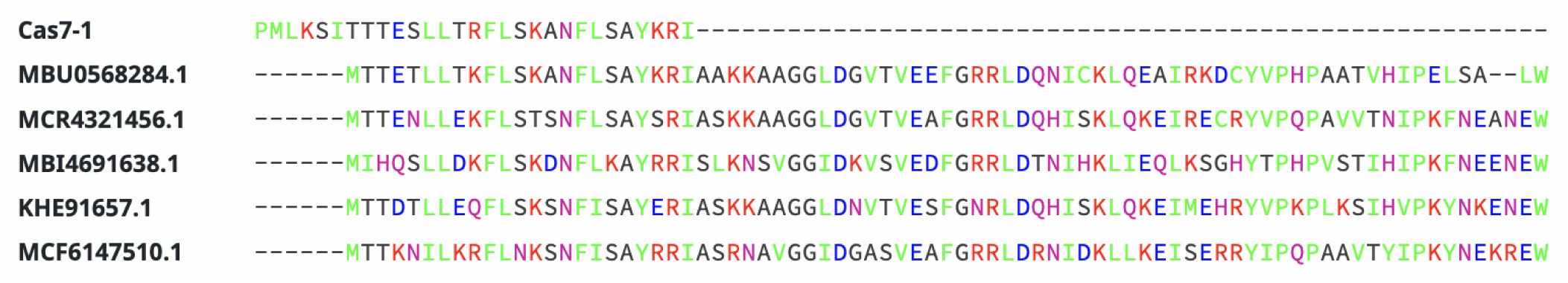
Alignment of 29AA N-terminus of Cas1 domain to RT-Cas1 fusion proteins. The Cas7-1 Cas1 N-terminal linker domain aligns well with multiple N-terminal regions of RT-Cas1 fusion proteins. NCBI ascension identities listed for each RT-Cas1 protein.

**Figure S6.**
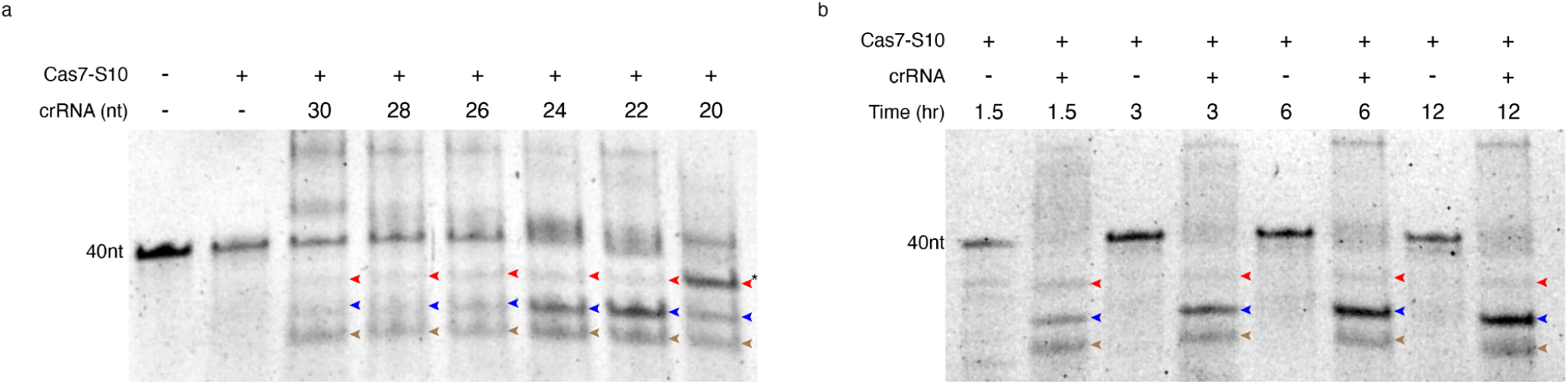
Biochemical assessment of Cas7-S10. (A) Examination of the influence of spacer length on cleavage activity of Cas7-S10. Red arrow indicates cleavage products from Cas7.2 domain, blue arrow indicates cleavage products from Cas7.3 and/or Cas7.2/Cas7.3 domain(s), and brown arrow indicates 20nt product from DR processing. The asterisk next to the 20nt spacer Cas7.2 cleavage indicates a cleavage product and processed crRNA. (B) Assessment of cleavage activity for Cas7-S10 over time (1.5 hr, 3 hr, 6 hr, and 12 hr).

**Figure S7.**
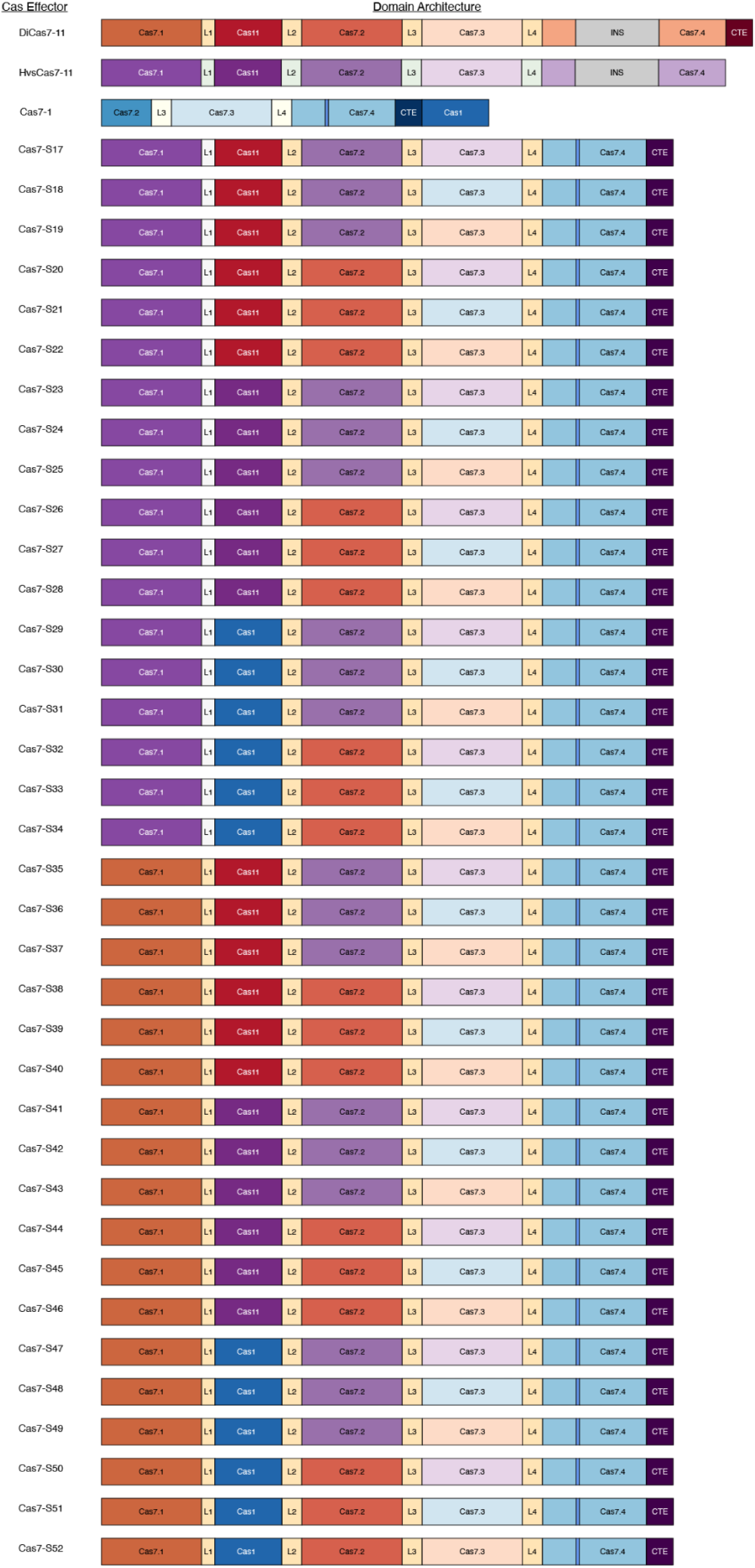
Domain breakdown of compact Cas7-S variants. Original proteins to build the initial Cas7-S variants are DiCas7-11 (red), HvsCas7-11 (purple), and Cas7-1 (blue). Color scheme represents which native protein individual domains originate

**Figure S8.**
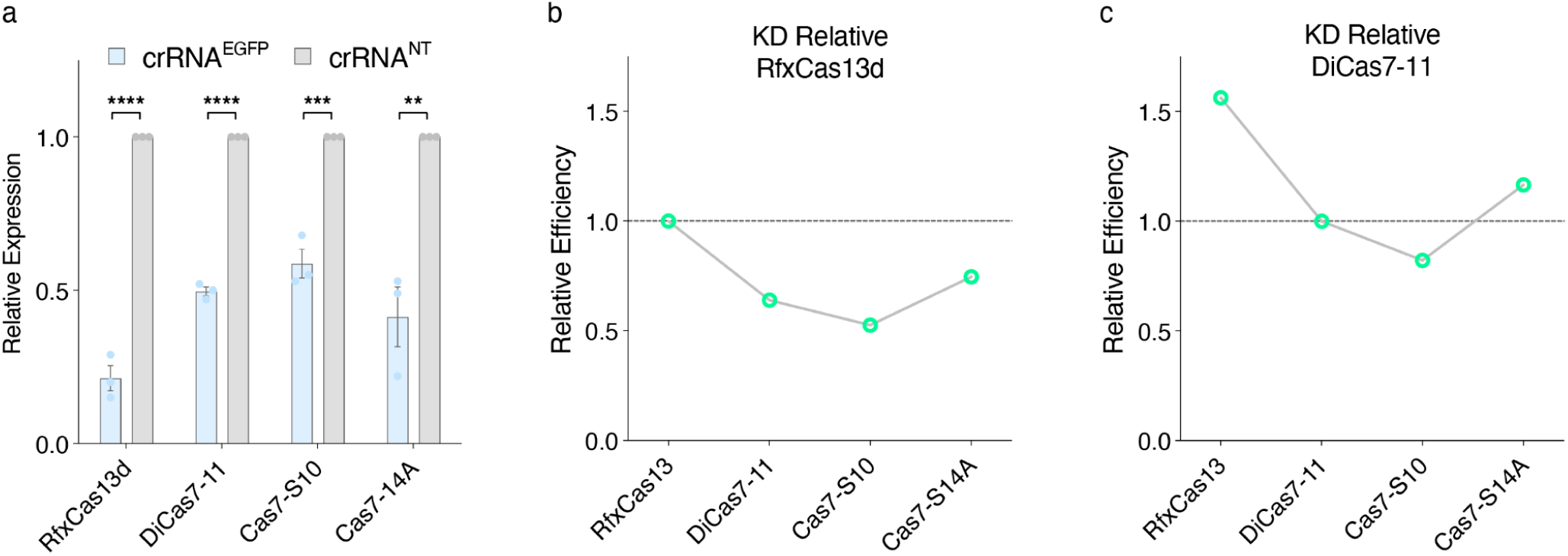
Cas7-S Targeted RNA knockdown comparison against RfxCas13d and DiCas7-11. (**A**) qPCR analysis of RNA knockdown targeting GFP in HEK293T cells. (**B**) GFP knockdown of each effector relative to RfxCas13d. (**C**) GFP knockdown of each effector relative to DiCas7-11. Significance is calculated and determined using unpaired t-test between crRNA^EGFP^ and crRNA^NT^. Error bars represent sem (n = 3).

**Figure S9.**
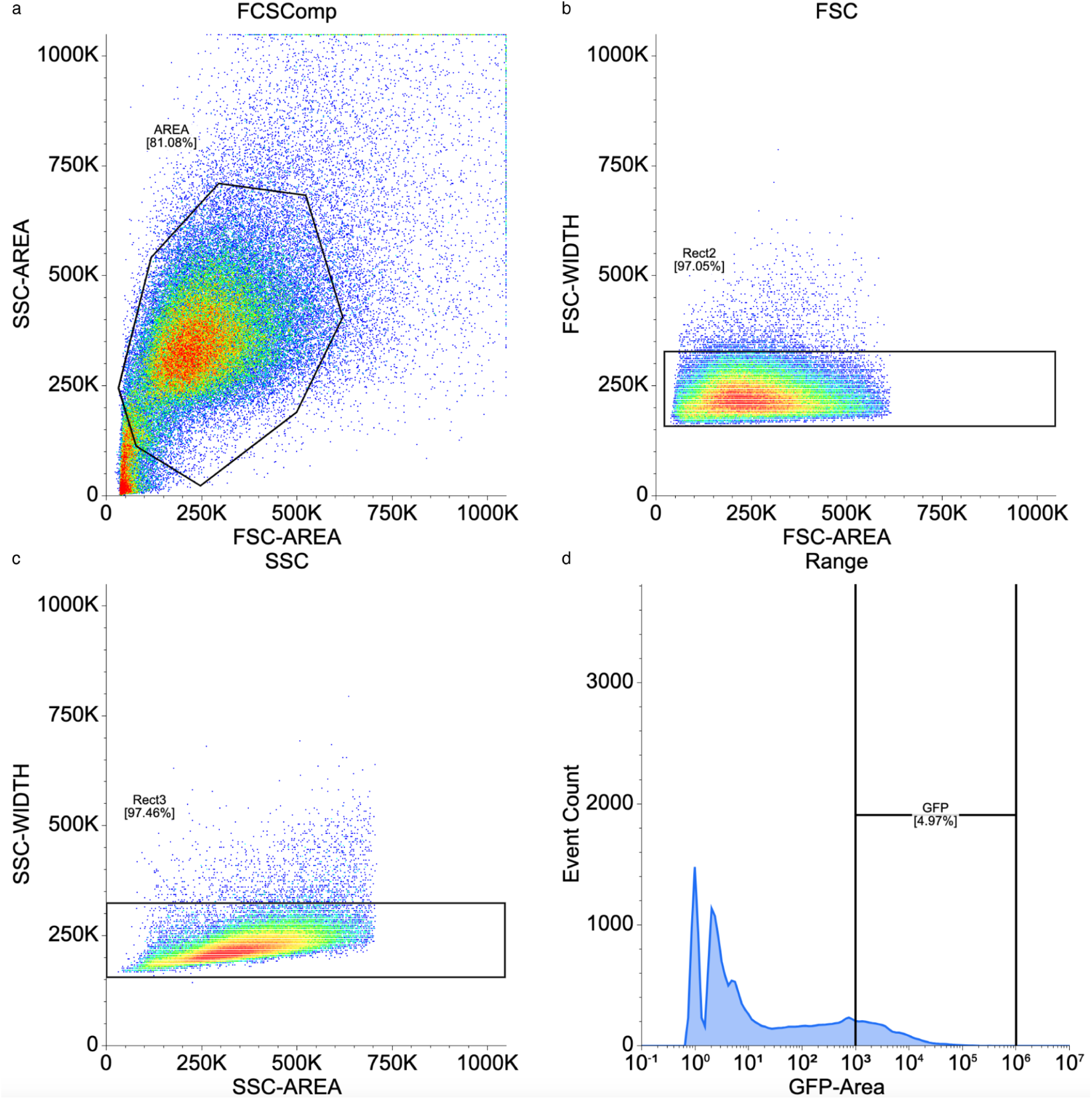
Representative gating method for flow cytometry analysis of EGFP knockdown. (**A**) Area of cells with polygon identifying selected cells. (**B**) Further refinement of selected cells – FSC width vs area. (**C**) Further refinement of selected cells – SSC width vs area. (**D**) Defined range of high EGFP population.

## References

1. K. S. Makarova, Y. I. Wolf, J. Iranzo, S. A. Shmakov, O. S. Alkhnbashi, S. J. J. Brouns, E. Charpentier, D. Cheng, D. H. Haft, P. Horvath, S. Moineau, F. J. M. Mojica, D. Scott, S. A. Shah, V. Siksnys, M. P. Terns, Č. Venclovas, M. F. White, A. F. Yakunin, W. Yan, F. Zhang, R. A. Garrett, R. Backofen, J. van der Oost, R. Barrangou, E. V. Koonin, Evolutionary classification of CRISPR-Cas systems: a burst of class 2 and derived variants. Nat. Rev. Microbiol. 18, 67–83 (2020).

2. S. P. B. van Beljouw, A. C. Haagsma, A. Rodríguez-Molina, D. F. van den Berg, J. N. A. Vink, S. J. J. Brouns, The gRAMP CRISPR-Cas effector is an RNA endonuclease complexed with a caspase-like peptidase. Science. 373, 1349–1353 (2021).

3. A. Özcan, R. Krajeski, E. Ioannidi, B. Lee, A. Gardner, K. S. Makarova, E. V. Koonin, O. O. Abudayyeh, J. S. Gootenberg, Programmable RNA targeting with the single-protein CRISPR effector Cas7-11. Nature, 1–6 (2021).

4. K. Kato, W. Zhou, S. Okazaki, Y. Isayama, T. Nishizawa, J. S. Gootenberg, O. O. Abudayyeh, H. Nishimasu, Structure and engineering of the type III-E CRISPR-Cas7-11 effector complex. Cell. 0 (2022), doi:10.1016/j.cell.2022.05.003.

5. H. N. Goswami, J. Rai, A. Das, H. Li, Molecular mechanism of active Cas7-11 in processing CRISPR RNA and interfering target RNA. Elife. 11 (2022), doi:10.7554/eLife.81678.

6. C. Hu, S. P. B. van Beljouw, K. H. Nam, G. Schuler, F. Ding, Y. Cui, A. Rodríguez-Molina, A. C. Haagsma, M. Valk, M. Pabst, S. J. J. Brouns, A. Ke, Craspase is a CRISPR RNA-guided, RNA-activated protease. Science. 377, 1278–1285 (2022).

7. A. B. Buchman, D. J. Brogan, R. Sun, T. Yang, P. D. Hsu, O. S. Akbari, Programmable RNA Targeting Using CasRx in Flies. CRISPR J. 3, 164–176 (2020).

8. J. F. Bot, J. van der Oost, N. Geijsen, The double life of CRISPR-Cas13. Curr. Opin. Biotechnol. 78, 102789 (2022).

9. C. P. Kelley, M. C. Haerle, E. T. Wang, Negative autoregulation mitigates collateral RNase activity of repeat-targeting CRISPR-Cas13d in mammalian cells. Cell Rep. 40, 111226 (2022).

10. Q. Wang, X. Liu, J. Zhou, C. Yang, G. Wang, Y. Tan, Y. Wu, S. Zhang, K. Yi, C. Kang, The CRISPR-Cas13a Gene-Editing System Induces Collateral Cleavage of RNA in Glioma Cells. Adv. Sci. 6, 1901299 (2019).

11. K. Kato, S. Okazaki, C. Schmitt-Ulms, K. Jiang, W. Zhou, J. Ishikawa, Y. Isayama, S. Adachi, T. Nishizawa, K. S. Makarova, E. V. Koonin, O. O. Abudayyeh, J. S. Gootenberg, H. Nishimasu, RNA-triggered protein cleavage and cell growth arrest by the type III-E CRISPR nuclease-protease. Science. 378, 882–889 (2022).

12. B. Ekundayo, D. Torre, B. Beckert, S. Nazarov, A. Myasnikov, H. Stahlberg, D. Ni, Structural insights into the regulation of Cas7-11 by TPR-CHAT. Nat. Struct. Mol. Biol. 30, 135–139 (2023).

13. K. S. Makarova, Y. I. Wolf, O. S. Alkhnbashi, F. Costa, S. A. Shah, S. J. Saunders, R. Barrangou, S. J. J. Brouns, E. Charpentier, D. H. Haft, P. Horvath, S. Moineau, F. J. M. Mojica, R. M. Terns, M. P. Terns, M. F. White, A. F. Yakunin, R. A. Garrett, J. van der Oost, R. Backofen, E. V. Koonin, An updated evolutionary classification of CRISPR-Cas systems. Nat. Rev. Microbiol. 13, 722–736 (2015).

14. W. X. Yan, S. Chong, H. Zhang, K. S. Makarova, E. V. Koonin, D. R. Cheng, D. A. Scott, Cas13d Is a Compact RNA-Targeting Type VI CRISPR Effector Positively Modulated by a WYL-Domain-Containing Accessory Protein. Mol. Cell. 70, 327–339.e5 (2018).

15. S. Konermann, P. Lotfy, N. J. Brideau, J. Oki, M. N. Shokhirev, P. D. Hsu, Transcriptome Engineering with RNA-Targeting Type VI-D CRISPR Effectors. Cell. 173, 665–676.e14 (2018).

16. S. Silas, G. Mohr, D. J. Sidote, L. M. Markham, A. Sanchez-Amat, D. Bhaya, A. M. Lambowitz, A. Z. Fire, Direct CRISPR spacer acquisition from RNA by a natural reverse transcriptase–Cas1 fusion protein. Science. 351 (2016), , doi:10.1126/science.aad4234.

17. J. Jumper, R. Evans, A. Pritzel, T. Green, M. Figurnov, O. Ronneberger, K. Tunyasuvunakool, R. Bates, A. Žídek, A. Potapenko, A. Bridgland, C. Meyer, S. A. A. Kohl, A. J. Ballard, A. Cowie, B. Romera-Paredes, S. Nikolov, R. Jain, J. Adler, T. Back, S. Petersen, D. Reiman, E. Clancy, M. Zielinski, M. Steinegger, M. Pacholska, T. Berghammer, S. Bodenstein, D. Silver, O. Vinyals, A. W. Senior, K. Kavukcuoglu, P. Kohli, D. Hassabis, Highly accurate protein structure prediction with AlphaFold. Nature. 596, 583–589 (2021).

18. M. Varadi, S. Anyango, M. Deshpande, S. Nair, C. Natassia, G. Yordanova, D. Yuan, O. Stroe, G. Wood, A. Laydon, A. Žídek, T. Green, K. Tunyasuvunakool, S. Petersen, J. Jumper, E. Clancy, R. Green, A. Vora, M. Lutfi, M. Figurnov, A. Cowie, N. Hobbs, P. Kohli, G. Kleywegt, E. Birney, D. Hassabis, S. Velankar, AlphaFold Protein Structure Database: massively expanding the structural coverage of protein-sequence space with high-accuracy models. Nucleic Acids Res. 50, D439–D444 (2022).

19. M. A. Cianfrocco, M. Wong-Barnum, C. Youn, R. Wagner, A. Leschziner, “COSMIC2: A Science Gateway for Cryo-Electron Microscopy Structure Determination” in Proceedings of the Practice and Experience in Advanced Research Computing 2017 on Sustainability, Success and Impact (Association for Computing Machinery, New York, NY, USA, 2017; 10.1145/3093338.3093390), PEARC17, pp. 1–5.

20. Y. Zheng, J. Li, B. Wang, J. Han, Y. Hao, S. Wang, X. Ma, S. Yang, L. Ma, L. Yi, W. Peng, Endogenous Type I CRISPR-Cas: From Foreign DNA Defense to Prokaryotic Engineering. Front Bioeng Biotechnol. 8, 62 (2020).

21. D. J. Brogan, D. Chaverra-Rodriguez, C. P. Lin, A. L. Smidler, T. Yang, L. M. Alcantara, I. Antoshechkin, J. Liu, R. R. Raban, P. Belda-Ferre, R. Knight, E. A. Komives, O. S. Akbari, Development of a Rapid and Sensitive CasRx-Based Diagnostic Assay for SARS-CoV-2. ACS Sens (2021), doi:10.1021/acssensors.1c01088.

22. A. East-Seletsky, M. R. O’Connell, S. C. Knight, D. Burstein, J. H. D. Cate, R. Tjian, J. A. Doudna, Two distinct RNase activities of CRISPR-C2c2 enable guide-RNA processing and RNA detection. Nature. 538, 270–273 (2016).

23. B. Zetsche, J. S. Gootenberg, O. O. Abudayyeh, I. M. Slaymaker, K. S. Makarova, P. Essletzbichler, S. E. Volz, J. Joung, J. van der Oost, A. Regev, E. V. Koonin, F. Zhang, Cpf1 is a single RNA-guided endonuclease of a class 2 CRISPR-Cas system. Cell. 163, 759–771 (2015).

24. K. K. Yang, Z. Wu, F. H. Arnold, Machine-learning-guided directed evolution for protein engineering. Nat. Methods. 16, 687–694 (2019).

25. I. S. Al’Abri, D. J. Haller, Z. Li, N. Crook, Inducible directed evolution of complex phenotypes in bacteria. Nucleic Acids Res. 50, e58 (2022).

26. F. Adeola, Normalization of Gene Expression by Quantitative RT-PCR in Human Cell Line: comparison of 12 Endogenous Reference Genes. Ethiop. J. Health Sci. 28, 741–748 (2018).

